# High regional and intra-generic variation in susceptibility to mass bleaching in Indo-Pacific coral species

**DOI:** 10.1101/2021.01.10.426149

**Authors:** Paul R. Muir, Terence Done, J. David Aguirre

## Abstract

**Aim:** Mass bleaching is a major threat to reef-building corals and the ecosystems they underpin. Here, we identified regional variation in the nature of this threat in terms of the bleaching-susceptibility of individual coral species on some Indian Ocean and Pacific Ocean reefs.

**Location:** 22 sites in the central Great Barrier Reef, Australia (GBR) and 30 sites in the central Maldives Archipelago (MA).

**Time period:** 2002 for the GBR and 2016 for the MA.

**Major taxa studied:** Corals (Order Scleractinia).

**Methods:** Following marine heat-wave conditions, timed in-situ surveys were used to record bleaching responses (tissue colour) of large samples of individual coral colonies. Responses of 106 shared species were analysed for sites with similar levels of temperature stress, depth of occurrence and mortality. In each region, phylogenetic mixed models were used to partition the effects on responses of species of deep-time phylogeny, contemporary history and local-scale, among-site variability.

**Results:** Relative susceptibility to bleaching varied widely between regions: only 27 of the 106 shared species were in the same quartile for relative susceptibility in both regions. Few species were highly susceptible in both regions. Closely related species varied widely in their individual susceptibilities. Phylogenetic effects were moderate in both regions, but contemporary phenotypic effects indicative of recent evolution and acclimatization were greater in the MA, consistent with a stronger history of recent bleaching.

**Main conclusions:** The high regional and intra-generic variation in coral bleaching-susceptibility described here suggests there may be important differences in the extent to which these Indian and Pacific Ocean coral populations are exhibiting responses to deep-time evolutionary changes on the one hand, versus recent adaptation, on the other. There is a concerning scarcity of this type of data, by which coral species most at risk from bleaching in particular regions may be more accurately identified.

## 1. INTRODUCTION

Coral reefs are one of the most threatened ecosystems on the planet, with mass coral bleaching widely regarded as their greatest current threat (Heron et al., 2016). Repeated, severe bleaching events have significantly damaged reefs across the tropics and subtropics, and mass bleaching events are predicted to increase in frequency and severity in the near future (Hoegh-Guldberg et al., 2007; Hughes et al., 2017; Hughes, et al., 2018a). Indeed, in 2020, the Great Barrier Reef (GBR) underwent its third severe event within three years (Hughes & Pratchett, 2020). Local and possible regional species extinctions related to repeated bleaching events have been reported (Glynn, 2011; Muir et al., 2017; Sheppard et al., 2020) and there is now a risk of global extinctions of reef-building coral species (Carpenter et al., 2008; IUCN, 2017; Richards & Day, 2018).

Rising sea temperatures, the ultimate cause of most mass bleaching events (Brown, 1997), are a global phenomenon, but the search for pragmatic management interventions are focused at local and regional scales (Anthony et al., 2015; Wooldridge & Done, 2009). While conventional management measures continue in the form of reducing impacts from agricultural runoff, overfishing and coastal development (GBRMPA, 2018; Steneck et al., 2019), current biological research seeks to support active interventions such as reef restoration, assisted evolution, assisted migration, hybridisation, seed banking, artificial refugia and assisted recruitment (van Oppen et al., 2017). However, given the scale of the problem, with around 750 species of scleractinian reef-corals (Hoeksema & Cairns, 2019) and over 280,000 km^2^ of reefs threatened around the globe (Spalding et al., 2001), these management and research efforts face a daunting task.

Despite the threat to reef-building corals globally, there is surprisingly little data on the bleaching susceptibility of individual species. Susceptibility has been documented mostly for coral genera (Baird et al., 2018; Guest et al., 2012; Hughes et al., 2018b; McClanahan et al., 2007; Marshall & Baird, 2000; Chou et al., 2016; Loya et al., 2001) with only a small number of species examined. While wide geographical variation in relative susceptibility has been established for entire coral communities (McClanahan et al. 2020, Thompson & van Woesik, 2009), there are few comparative studies of individual taxa and these are mostly restricted to comparisons at a genus-level (Guest et al., 2012, McClanahan et al., 2004; Pratchett et al., 2013). While much of the current research is aimed at active interventions to prevent loss of coral species following repeated mass bleaching events, the species most at risk from bleaching have not been well documented. For instance, the IUCN “Red List” which is widely used to categorise the risk and conservation status of coral taxa, is compiled from mainly genus-level bleaching data from a small number of regions (Carpenter et al., 2008) and thus might be misrepresenting the status of many coral species and regional populations. The lack of data limits the extent to which management and interventions could be targeted to the most susceptible species.

Here, we investigate the susceptibility of a broad range of Indo-Pacific scleractinian species to moderate bleaching conditions in shallow reef waters of the central Maldives Archipelago (MA) during 2016 and the central Great Barrier Reef (GBR) during 2002. To the best of the authors’ knowledge, our datasets are unique in capturing in-situ bleaching responses for diverse assemblages of coral species. For each region, phylogenetic mixed model analyses were used to estimate longer-term phylogenetically heritable effects, shorter-term effects commensurate with adaptation and/or acclimatisation (hereafter, contemporary phenotypic effects) and local-scale variation. Comparison between regions were restricted to corals exposed to similar levels of temperature stress at a similar depth of occurrence, but since mass bleaching is caused by a complex array of stressors that can occur over many weeks (Skirving et al. 2019) we avoided absolute comparisons. Instead, we compared the responses of the 106 species present in both regions in terms of their relative susceptibilities. We then assessed how the available data might be used to estimate species risk across wide geographic regions and closely related species. The findings have important implications for how coral bleaching, the greatest ongoing threat to these critically important, yet vulnerable species, is assessed and how research and management efforts might be prioritized.

## 2. MATERIALS AND METHODS

For the GBR in 2002 and the MA in 2016, sea temperatures exceeded bleaching thresholds over wide areas for several weeks, resulting in moderate mortalities over the ensuing months. To control for variation due to differing levels of temperate stress we restricted our analyses to sites with a Degree Heating Week (DHW) index of between 4 and 7 derived from satellite data (NOAA, 2017). This index is widely used to estimate the duration and severity of the temperature anomaly, with 4-7 DHW currently considered a moderate event (NOAA, 2017; Skirving, et al., 2019). Colony mortalities at selected sites reached 68% in the MA (Cowburn et al. 2019) and 86% in the GBR (Done et al. 2003). Since there can be a marked depth gradient in bleaching response (Muir et al., 2017; Baird et al., 2018), we focussed on corals assayed between 3 to 11 m depth in both regions. We further confined our analyses to the proportion of colonies with severe bleaching or recently partial or complete mortality to account for the possibility that less affected individuals recovered before our surveys (see below).

### 2.1 Field Surveys

A total of 14 locations (22 sites) in the central GBR and 10 locations (30 sites) in the central MA met the DHW criteria for inclusion in the analyses (Figure 1). At each site, timed surveys using SCUBA or snorkel dives of 45-80 minutes duration were conducted, recording bleaching responses and recent mortality of individuals according to standard methods (Supporting Information, Table S1). Species identifications were mostly made *in situ*, with high-resolution macro photographs or small samples taken from problematic colonies (under permit) for later analysis by comparison with material available in the Queensland Museum, consultation with relevant experts and reference to standard texts (see Supporting Information for details). Both datasets were converted to current valid species (Hoeksema & Cairns, 2019). For certain genera (e.g. *Montipora, Porites, Goniopora* and *Cycloseris*) it was not possible to accurately identify many of the species *in situ*, so these were assessed as grouped species and only considered in a supplementary analysis.

**Figure 1.**
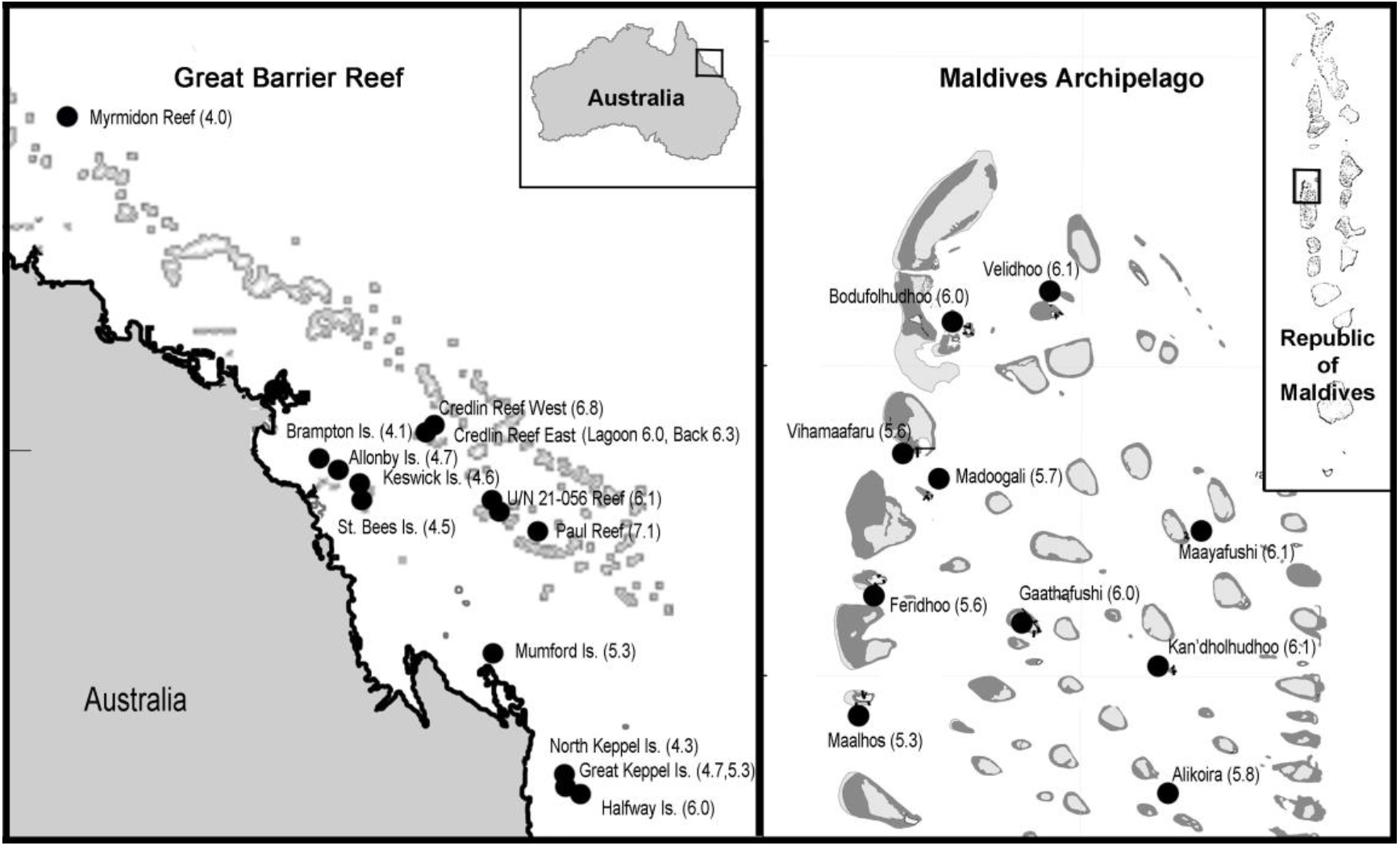
Locations where scleractinian coral bleaching responses were surveyed in the Great Barrier Reef and Maldives Archipelago. The severity of the bleaching at each location is given as peak degree heating weeks (DHW) in brackets.

The MA locations were surveyed two months after the peak of the temperature anomaly, whereas the GBR locations were surveyed 4-6 months after. To assess the effect of survey timing, we analysed a supplementary dataset of 142 individual corals (five species) monitored *in situ* after a moderate bleaching event in the GBR (Baird & Marshall, 2002, data kindly provided by A. Baird as: DOI to be supplied). These data showed that colonies with mild to moderate bleaching had reasonable recovery, whereas severely bleached colonies showed little recovery or >50% tissue death (Supporting Information, Table S2). Thus, the proportion of severely affected or recently dead/partially dead individuals remained consistent from 2.5 to 5.5 months after peak temperatures. We were therefore confident that our analyses were robust to differences in the timing of our surveys.

### 2.2 Statistical Analyses

The proportion of severely affected or recently dead/partially dead individuals were analysed using phylogenetic generalised linear mixed models implemented in the MCMCglmm package (Hadfield, 2019) in R v3.6.1. Separate models were applied to each region to accommodate different fixed effects. For the MA analysis, fixed effects included depth (3-5 m and 9-11 m), shading (shaded microhabitats and unshaded) and the depth × shading interaction. For the GBR analysis, the fixed effects included two levels of depth (3-5 m and 6-9 m), seven levels of habitat/zone (nearshore-fringing reef, coastal-fringing reef, offshore-northern lagoon, offshore-northern reef slope, offshore-southern back reef, offshore-southern lagoon and offshore-southern reef slope) and habitat/zone × depth interaction. The random effects used in both analyses were: phylogenetically heritable effects, contemporary phenotypic effects, and location. Only taxa with more than two records in each region were included in the analyses. See S2, Supporting Information for further detail.

The super tree of Huang and Roy (2016) was used in the phylogenetic mixed model analyses, it currently being the most comprehensive for the group. We also explored the effect of phylogenetic uncertainty on our results and undertook analyses using a restricted phylogeny based only on molecular data (Huang & Roy, 2016). These analyses are not considered further here as they recovered similar results to the analyses conducted using the super tree phylogeny, indicating our methods were robust to uncertainty in the construction of the super tree (Supporting Information, Tables S4 and S5). A supplementary analysis using the Bleaching Mortality Index (BMI, McClanahan et al., 2004) with species data pooled into genera was conducted to allow comparison with previous reports.

## 3. RESULTS

Analysis of 2888 individuals from 180 species, 55 genera and 14 families from the GBR showed moderate phylogenetically heritable effects, weak contemporary effects and weak local-scale effects (Table 1). For the MA, analysis of data from 4480 individuals representing 144 species, 47 genera and 14 families also showed moderate phylogenetic effects and minor local-scale effects. However, contemporary effects were 2.7 times greater than those for the GBR. In both regions, the species responses varied widely within genera (Figure 2). Genus *Acropora* had the largest number of species present in each region (35 species in both regions) and species were present in both the lowest and highest quartiles of bleaching response in each region (Figure 3). Species of the genera *Pocillopora, Echinopora, Leptastrea, Platygyra* and *Dipsastraea* showed a greater range of responses in the MA relative to the GBR (Figure 2).

**Figure 2.**
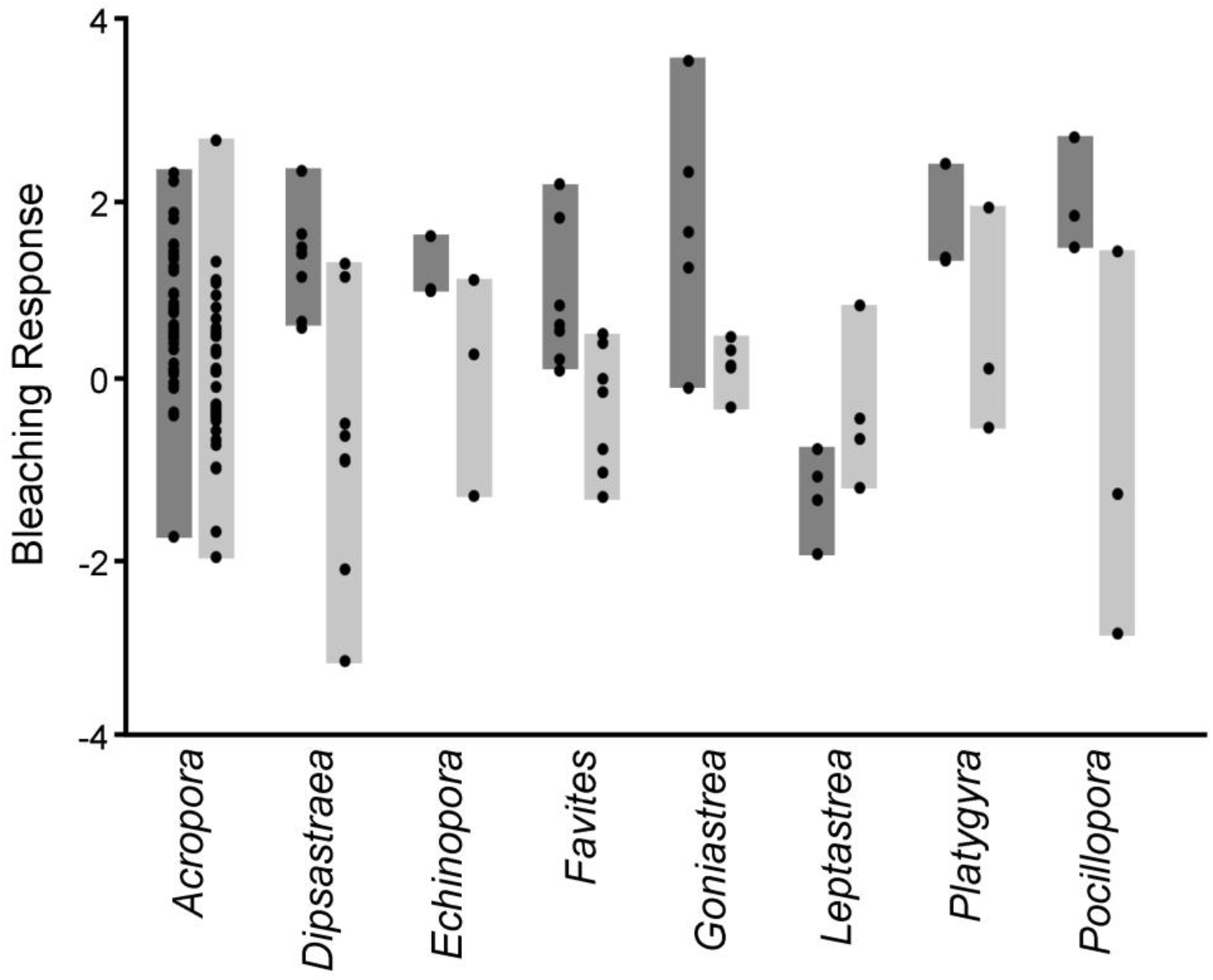
Species’ bleaching responses (●) varied widely within many of the main scleractinian coral genera in both the Great Barrier Reef 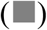 and Maldives Archipelago 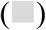. Responses shown as posterior medians derived from phylogenetic mixed model analyses. The genera shown had more than two shared species between regions with sufficient data for analysis.

**Figure 3.**
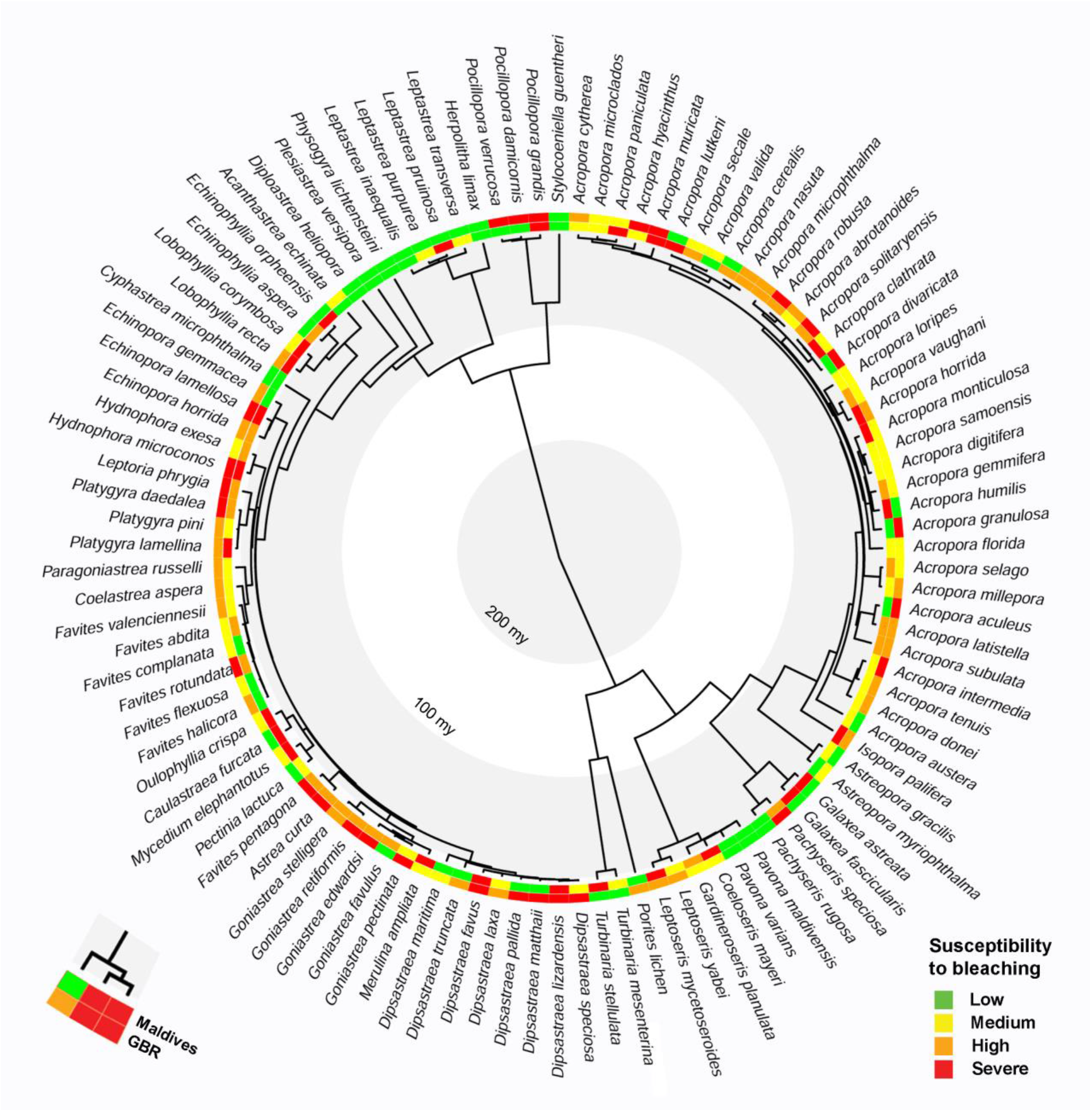
Susceptibility to moderate coral bleaching for scleractinian species in the Maldives Archipelago (inner ring) and Great Barrier Reef (outer ring) plotted onto the currently accepted phylogeny of the group (Huang and Roy, 2016). Susceptibility was categorized using quartiles of the species’ posterior medians derived from separate phylogenetic mixed models that analysed the proportion of the population that was severely affected.

**Table 1.** Summary of phylogenetic mixed model analyses for the bleaching responses of scleractinian species in the Maldives Archipelago and Great Barrier Reef (GBR). The proportion of variation in bleaching explained by phylogenetically heritable (Phylogenetic), recent adaptation/acclimatization (Contemporary) and local-scale effects. Proportions given as posterior medians with 0.025 and 0.975 quantiles in brackets.

| Region | Proportion of Variation Explained |  |  |
| --- | --- | --- | --- |
|  | Phylogenetic | Contemporary | Local-Scale |
| Maldives | 0.367 (0.065,0.638) | 0.115 (0.047, 0.258) | 0.032 (0.011,0.105) |
| GBR | 0.454 (0.264, 0.658) | 0.042 (0.094,0.103) | 0.025 (0.006,0.095) |

Of the 218 species with sufficient data for analysis, only 106 species (64 genera, 14 families) were present in both regions. Categorizing responses into quartiles based on their posterior median estimates from the phylogenetic mixed model analyses showed only 27 species (25.4%) were in the same quartile in both regions, 45 species differed by one quartile and 34 by two or more quartiles (Figure 3). Only six species were in the most susceptible quartile in both regions. The phylogenetically heritable effects for each species showed a weak correlation between regions, markedly different from a slope of +/-1.0 expected if bleaching responses were phylogenetically constrained between regions (Figure 4). The contemporary effects also showed a low correlation between regions (Figure 4).

**Figure 4.**
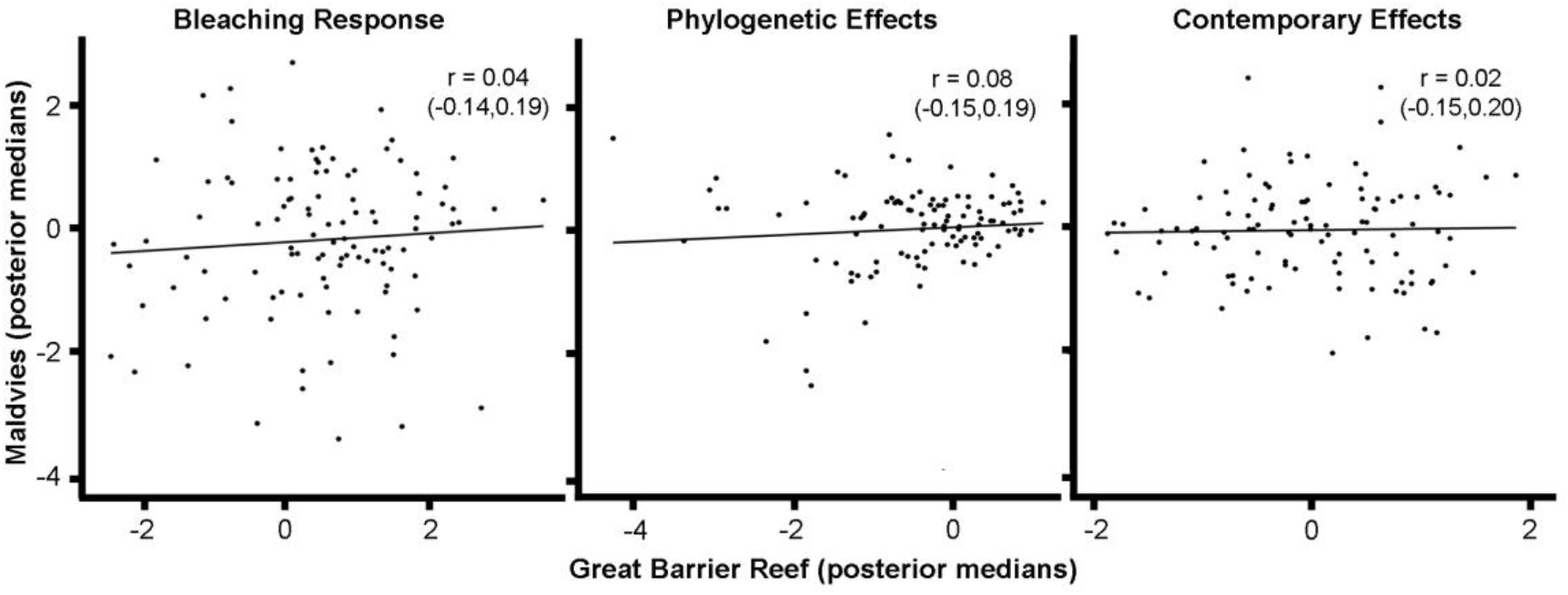
The bleaching responses of scleractinian species varied widely between regions, with phylogenetic mixed model analyses indicating that the variation had origins in both deep-time, phylogenetically heritable effects and short-term contemporary effects. These effects were poorly correlated between regions. Responses are shown as posterior medians (see Supporting Information, Figure S3 for details).

As contemporary effects were 2.7 times greater in the MA than the GBR (Table 1) and the region had a stronger history of recent bleaching, we examined species with the strongest contemporary effects consistent with recent adaptation or acclimatisation (“C” in Supporting Information, Table S3). A wide range of genera (21 of 47 analysed) were represented in the upper quartile (38 species) for contemporary effects in the MA, including the ecologically significant genera: *Acropora, Dipsastraea, Cycloseris, Favites, Goniastrea, Hydnophora, Lobophyllia, Pavona* and *Pocillopora*.

For many genera, re-analysis of our data using the bleaching mortality index (BMI) at the level of genus (Supporting Information, Figure S2) showed results somewhat similar to those of the study that defined it (McClanahan et al., 2004). However, the relative responses were markedly different to those derived from the phylogenetic mixed models of our main analyses.

## 4. DISCUSSION

We report the first regional comparison of bleaching susceptibility for a broad range of common Indo-Pacific reef corals at the level of species. Comparing shallow-reef populations exposed to similar levels of moderate thermal stress we found relative responses varied only slightly at local scales, but showed wide differences between the regions. The findings have important implications for how mass bleaching research is conducted and how this severe threat is addressed.

Prior to this study, the small amount of data available indicated low to moderate levels of regional variation in the relative susceptibility to mass bleaching for Indo-Pacific coral taxa. While entire assemblages varied widely in their overall susceptibility (McClanahan et al. 2020), the major coral genera often showed a similar hierarchy of susceptibility for different regions and bleaching events (e.g. Loya et al. 2001, Marshall & Baird 2000, Hughes et al. 2017). The few direct comparative studies also showed low to moderate regional variation. For example, relative bleaching responses between East Africa and the north/central GBR were similar for 19 coral genera (McClanahan et al., 2004), while a comparison of sites within SE Asia and Australia found only two of 18 genera varied significantly in their relative susceptibility for two sites (Guest et al., 2012). Our study showed markedly higher levels of regional variation when susceptibility was considered at the level of species. Only 25% of the 106 species with sufficient data for analysis were present in the same quartile of susceptibility in the MA and GBR (Figure 3). This comparison was for an identical assemblage of currently accepted species for similar levels of temperature stress and depth of occurrence and for events that produced similar levels of mortality.

Phylogenetic analyses allow the dependence resulting from a shared evolutionary history along a phylogeny to be considered (Housworth et al., 2004) and here provide an indication of the origins of variation in bleaching susceptibility between the two assemblages. A strong inter-regional correlation in the phylogenetic contribution to bleaching response would have suggested that bleaching susceptibility was strongly phylogenetically constrained. By contrast, we found only a weak correlation (Figure 4) suggesting divergence in deep-time that has affected susceptibility to bleaching in each region. This result, and indeed our general findings, are consistent with recent genetics studies that indicate wide divergence within many species of Indian and Pacific Ocean reef corals, fishes and other reef fauna (summarized Richards et al., 2016). Several cryptic regional endemic species or subspecies have also been resolved within species previously considered pan Indo-Pacific (Richards et al., 2016; Arrigoni et al., 2020), suggesting regional divergence of reef faunas may be greater than currently recognised. Such cryptic regional endemics would account for some the regional variation documented here. The existence of more regionally restricted taxa would have implications for management and conservation: taxa with smaller range sizes are likely to have a greater risk of extinction (Richards et al., 2016), further strengthening the case for species-level and regional assessments of susceptibility.

Our analyses indicate that phylogeny imposes only limited constraints on the bleaching responses in each region. This suggests that regional divergence of entire lineages, as well as recent acclimation and adaptation, play a role in mediating bleaching responses. The greater proportion of variation partitioned to contemporary effects in the MA assemblage in 2016, relative to the 2002 GBR assemblage is consistent with this view (Table 1). The MA study area experienced several bleaching events in the two decades before our surveys (summarized Muir et al., 2017), potentially providing a strong pressure for contemporary phenotypic responses. By contrast, although several events had been reported on the GBR in the two decades before our 2002 surveys (Berkelmans & Oliver, 1999; Berkelmans et al., 2004), our study sites had not yet been exposed to the globally widespread and severe events of the present century, and thus any associated selective tendency towards reduced susceptibility to subsequent bleaching.

Reduced susceptibility following a series of recent bleaching events has been reported previously, but mainly for communities (Thompson & van Woesik, 2009; McClanahan et al. 2020) or genera. Relatively low bleaching responses for the genera *Pocillopora* and *Acropora* in Singapore and Malaysia (Guest et al., 2012, Chou et al., 2016) and for *Acropora* and *Montipora* in French Polynesia (Pratchett et al., 2013) have been attributed to adaptation/acclimation following several bleaching events. Here, we partitioned individual species responses between deep-time and contemporary effects to better understand the potential for adaption and/or acclimation. This method could provide a means of selecting species for different streams of management and intervention. For example, in the MA the species with the strongest contemporary phenotypic resistance following repeated bleaching events (“C” in Supporting Information, Table S3) are potential candidates for interventions such as reef restoration (Chamberland et al., 2017; dela Cruz & Harrison, 2017), assisted migration (van Oppen et al., 2017) and isolation of genes that confer resistance (van Oppen et al., 2015). Conversely, species with weak contemporary effects and higher bleaching responses (“P” in Supporting Information, Table S3) are potentially at increased risk and require heightened monitoring and first consideration for interventions such as artificial refugia (Coelhoa et al., 2017), assisted evolution (van Oppen, et al., 2015) and ‘seed-banking’ of cryopreserved gametes or larvae (Daly et al., 2018). Clearly, more data from other events and regions are required to fully realize the potential offered by this approach: our categories are only applicable for the MA and will likely change as the populations are challenged by further bleaching events.

High regional and intra-generic variation in bleaching susceptibility have important implications for the prioritization of reef management and conservation. The IUCN Red List (Carpenter et al., 2008) is widely used to set these priorities, but is currently based on mostly genus-level bleaching data from few locations. Obtaining the data to address this issue may be challenging as many species require expert taxonomic input for accurate identification, the dominant genera *Acropora, Porites and Montipora* being particularly difficult. In addition, high local-scale variation can be problematic for *in situ* bleaching assessments (Chou et al., 2016; Cantin & Spalding, 2008), but our methods produced low local-scale variation and consistent species responses within each region (Table 1). Thus, despite the challenges there are the means to better inform management priorities and mitigate species loss.

By assessing coral bleaching responses at greater taxonomic resolution over a wide range of taxa, we found much greater variation within genera and between regions than had previously been reported. By analysing these data using phylogenetic comparisons we found evidence for this variation having origins in both deep-time and recent adaptation and/or acclimation in response to repeated events. Overall, we conclude that there is an alarming lack of reliable data on species’ susceptibility with which to address the coral bleaching phenomenon, the greatest current threat to these already endangered species. Somewhat encouragingly, we found that only a small proportion of species were highly susceptible in both regions and that bleaching responses were weakly phylogenetically constrained, with some potential for adaptation and/or acclimation to moderate levels of bleaching stress.

## Acknowledgements

We thank Paul Marshall, Ameer Abdullah, Hussein Zahir, Emre Turak and Mary Wakeford for assisting with fieldwork and Michel Pichon and Carden Wallace for taxonomic advice to both teams. Emre Turak provided much of the identifications for the GBR surveys. Ray Berkelmans, Tom Schlesinger, Michel Pichon and Cynthia Riginos provided valuable advice and Andrew Baird kindly provided the supplementary bleaching recovery data.

