## Supplementary material for "High regional and intra-generic variation in susceptibility to mass bleaching in Indo-Pacific coral species"

### Identifications

The majority of identification of corals was *in situ* by experts in the para-taxonomy of the region, but for certain morpho-types high resolution photographs and/or small samples were taken under permit. Samples were tagged, bleached in hypochlorite solution for 24-48 h, rinsed in freshwater, dried and later examined in the laboratory using a stereo-microscope. Identification was made with reference to standard texts (Veron, 1986, 2000; Wallace, 1999; Wallace et al., 2012, Veron & Pichon, 1976) and in consultation with taxonomic experts M. Pichon and C. Wallace, Queensland Museum (QM). The extensive collections of the QM were also used for reference throughout the two studies.

### Analytical Methods

The common assumption that genus-level bleaching assessments are representative of the species-level responses, exemplifies a phylogenetic perspective – the bleaching responses of closely related individuals are more similar than the bleaching responses of more distantly related individuals. Accordingly, we might consider estimating phylogenetic effects as an extension of the current ideology which acknowledges that the phylogenetic distance between individuals and their most recent common ancestor is not uniform. Importantly, the relative magnitude of the phylogenetic contribution to phenotypic variation provides an indication of the contribution of macro-evolutionary and biogeographic processes to similarities and dissimilarities in the bleaching responses of individuals. Furthermore, if individual-level data are collected, then the effects of contemporary phenotypic effects such as adaptation and acclimatisation on bleaching can be better understood.

Analyses were implemented specifying the distribution of the response variable as a binary variable and with a logit link function (de Villemereuil & Nakagawa, 2014; Hadfield, 2016). A value of 1 for the response indexed severely bleached/recently dead individuals and a value of 0 indexed moderately bleached/unaffected individuals. For both regional models, we assumed a Cauchy prior for the fixed effects with mean of zero and a variance of $\sqrt{\pi^2/3}$. For the random effects, we assumed a Cauchy distribution with a mean of zero and variance equal to the observed variance in bleaching. For the residual term we fixed the prior at one, which ensured that the absolute value of the latent variable did not exceed twenty (Hadfield & Nakagawa, 2010). We stored 1000 posterior samples of the distribution of each parameter by specifying a total of 1,030,000 iterations, with a burn-in of 30,000 iterations and a thinning interval of 1000 iterations. We confirmed that the models were a good fit for the data by ensuring that the posterior predictive distribution of the median bleaching overlapped the observed median bleaching. Moreover, we confirmed that our results were robust to prior specification by adjusting the prior values to unreasonably large and small values, then examining changes in the posterior distributions of the parameter estimates.

For the phylogenetic component, we calculated the phylogenetic numerator relationship matrix (Hadfield & Nakagawa, 2010), based on the subtree of the scleractinian coral super-tree (Huang & Roy, 2016) that retained the combined species lists for both regions. Accordingly, because the same subtree was used for both datasets, the same phylogenetic numerator relationship matrix was used for both datasets, allowing fair comparisons of the magnitude of the phylogenetic effects between regions. To examine the robustness of our results to uncertainty in the construction of the phylogeny, we repeated each model for the 1000 samples of the posterior distribution of the super-tree. The results we present were for the tree with the smallest squared Robinson-Fould's distance to all possible trees, i.e. the median tree. This was tree number 663 of the posterior distribution of trees presented in the supplementary materials of Huang & Roy (2016).

To examine differences in the relative contribution of phylogenetic effects (i.e. the phylogenetic signal or phylogenetic heritability), and the species-level effects to variation in bleaching susceptibility in each region, we calculated the intra-class correlations for each sample of the posterior distributions of the respective random effects. The posterior distribution of the between-region correlation was then calculated for each sample of the posterior distributions of the phylogenetic, contemporary and the total species' susceptibility (calculated as the sum of the phylogenetic and contemporary components). Although correlations are bound to between -1 and 1, and therefore the posterior distribution could not overlap ±1, visual inspection of each effect suggested there was little support for the correlations overlapping ±1, and further analysis was considered unnecessary.

To explore the influence of uncertainty in our reported summary statistics as a result of the phylogeny used, we conducted two additional checks. First, because the placement of the tips and internal nodes within the phylogeny differs among posterior samples of the super-tree (Huang & Roy, 2016), we repeated our phylogenetic linear mixed model analyses for all samples of the posterior distribution to account for uncertainty in the construction of the phylogeny (Table S4). Second, as molecular data are considered more reliable for constructing phylogenies for this group, we also conducted a set of analyses (Table S5) using only the molecular phylogeny presented in (Huang & Roy, 2016). The molecular only phylogeny captured 155 of the species included in the super-tree phylogeny. Finally, to examine the effects of excluding groups of difficult to identify species from our main analyses (e.g. massive *Porites*, encrusting *Porites, Goniopora* species, *Montipora* species), we conducted a further analysis with these groups included (Table S5).

| **Maldives** | **GBR** | **For Analyses** |
|---|---|---|
| Unaffected (0) | Unaffected (1) | |
| Slightly pale (1) | Slightly pale (2) | |
| < 25% white (2) | | Mild or unaffected (0) |
| 25-50% white (3) | Partly white(3) | |
| 50-75% white (4) | | |
| >75% white (5) | White (4) | Severely bleached/ very recent or partially mortality (1) |
| Very recent or partial mortality (6) | Very recent or partial mortality (5) | |

**Table S1** Bleaching categories used for individual colonies during surveys and recoded for statistical analyses.

| | Months Since Peak Temperatures | | | | |
|---|---|---|---|---|---|
| **Species** | **2.5** | **3.5** | **4.5** | **5.5** | **N** |
| *Acropora millepora* | 0.25/0.48 | 0.25/0.18 | 0.25/0.10 | 0.25/0.03 | 40 |
| *Acropora hyacinthus* | 0.66/0.27 | 0.60/0.13 | 0.60/0.00 | 0.60/0.00 | 30 |
| *Seriatopora hystrix* | 0.11/0.81 | 0.07/0.81 | 0.07/0.41 | 0.07/0.00 | 27 |
| *Platygyra. daedalea** | 0.40/0.60 | 0.10/0.77 | 0.07/0.77 | 0.07/0.53 | 30 |
| *Porites spp.* | 0.00/1.00 | 0.00/0.93 | 0.00/0.73 | 0.00/0.07 | 15 |
| **Overall** | **0.32/0.58** | **0.23/0.49** | **0.22/0.35** | **0.23/0.13** | **142** |

**Table S2** The proportion of individuals reported as 100% bleached or dead for each month following a moderate mass bleaching event in the central GBR during 1998 (Baird & Marshall, 2002). The proportion of colonies with mild to moderate bleaching (categories from study 2-4) is given after the backslash. Individuals remained undisturbed *in situ* though the study period. *The massive coral *P. daedalea* was the only species to show apparent recovery from severe bleaching (i.e. in tissue colour), but was severely affected in terms of partial mortality. After 5.5 months 44% of colonies had >50% tissue death by area (Baird & Marshall, 2002). Original data kindly supplied by A. Baird, DOI: to be supplied.

| | | Total | | Phylogen. | | Contemp. | | | |
|---|---|---|---|---|---|---|---|---|---|
| **No** | **Species** | **GBR** | **RM** | **GBR** | **RM** | **GBR** | **RM** | **Region** | **Cat.** |
| 1 | *Acanthastrea echinata* | 0.24 | -2.29 | -1.26 | -0.83 | -0.05 | 0.18 | Both | - |
| 2 | *Acanthastrea hemprichii* | -1.29 | - | -1.39 | - | -0.96 | - | GBR | - |
| 3 | *Acanthastrea pachysepta* | -1.79 | - | -1.65 | - | -1.80 | - | GBR | - |
| 4 | *Acropora abrotanoides* | 0.85 | -0.43 | -0.11 | 0.00 | 0.81 | 0.54 | Both | C |
| 5 | *Acropora aculeus* | 1.50 | -2.03 | 0.35 | -0.14 | 1.09 | -0.86 | Both | - |
| 6 | *Acropora acuminata* | - | 0.07 | - | -0.32 | - | 0.16 | RM | - |
| 7 | *Acropora anthocercis* | 0.25 | - | -0.56 | - | -1.13 | - | GBR | - |
| 8 | *Acropora aspera* | 1.49 | - | -0.37 | - | -0.08 | - | GBR | - |
| 9 | *Acropora austera* | -0.42 | -0.70 | -0.41 | -0.91 | -1.39 | -0.20 | Both | - |
| 10 | *Acropora caroliniana* | - | 0.69 | - | -0.62 | - | -1.05 | RM | P |
| 11 | *Acropora cerealis* | -0.38 | 0.08 | -0.65 | 0.43 | -1.09 | -0.02 | Both | - |
| 12 | *Acropora clathrata* | 0.52 | 1.31 | 0.76 | 0.27 | 0.60 | 0.50 | Both | C |
| 13 | *Acropora cytherea* | 1.35 | -0.37 | -0.38 | 0.00 | 0.57 | 0.34 | Both | - |
| 14 | *Acropora digitifera* | 0.17 | -0.40 | -0.82 | 0.45 | -1.80 | 0.11 | Both | - |
| 15 | *Acropora divaricata* | 1.39 | -1.02 | 0.83 | -0.08 | -0.72 | -0.87 | Both | - |
| 16 | *Acropora donei* | 1.25 | -0.35 | 0.28 | -0.24 | 0.09 | -0.19 | Both | - |
| 17 | *Acropora elegans* | - | -3.07 | - | -0.27 | - | -1.17 | RM | - |
| 18 | *Acropora elseyi* | - | 0.30 | - | 0.25 | - | 0.30 | RM | - |
| 19 | *Acropora florida* | 0.52 | -0.42 | -0.35 | -0.35 | -1.78 | -0.36 | Both | - |
| 20 | *Acropora gemmifera* | 0.32 | 0.32 | -0.08 | 0.14 | 1.16 | 0.60 | Both | C |
| 21 | *Acropora glauca* | 1.64 | - | 0.01 | - | 0.23 | - | GBR | - |
| 22 | *Acropora grandis* | 0.49 | - | -0.88 | - | 0.26 | - | GBR | - |
| 23 | *Acropora granulosa* | 1.51 | -1.74 | -0.23 | 0.32 | -0.82 | -1.27 | Both | - |
| 24 | *Acropora horrida* | 0.57 | 0.93 | 0.08 | 0.05 | -0.49 | 0.08 | Both | - |
| 25 | *Acropora humilis* | -0.11 | 0.80 | 0.35 | 0.18 | -0.20 | 1.22 | Both | C |
| 26 | *Acropora hyacinthus* | 1.43 | -0.31 | -0.50 | 0.53 | 0.91 | -0.68 | Both | - |
| 27 | *Acropora intermedia* | 1.80 | -0.76 | 0.13 | -0.52 | -0.09 | 0.45 | Both | - |
| 28 | *Acropora kirstyae* | - | 0.91 | - | 0.63 | - | -0.25 | RM | - |
| 29 | *Acropora kosurini* | - | -0.95 | - | -0.32 | - | -0.67 | RM | - |
| 30 | *Acropora latistella* | 1.21 | 0.27 | 0.48 | 0.53 | -0.01 | 0.01 | Both | - |
| 31 | *Acropora listeri* | 1.27 | - | -0.79 | - | -0.21 | - | GBR | - |
| 32 | *Acropora longicyathus* | 1.66 | - | 0.61 | - | 0.49 | - | GBR | - |
| 33 | *Acropora loripes* | 0.46 | -0.48 | 0.04 | -0.26 | -0.72 | -0.74 | Both | - |
| 34 | *Acropora lutkeni* | -1.79 | 1.11 | -1.11 | 0.23 | 0.49 | 0.90 | Both | C |
| 35 | *Acropora microclados* | 0.09 | -0.41 | -0.09 | 0.48 | -0.21 | -0.04 | Both | - |

| | | | | | | | | | |
|---|---|---|---|---|---|---|---|---|---|
| 36 | *Acropora microphthalma* | 0.60 | 0.07 | -0.52 | 0.31 | 1.26 | 0.56 | Both | C |
| 37 | *Acropora millepora* | 0.76 | -0.59 | -0.64 | -0.38 | -0.24 | -0.51 | Both | - |
| 38 | *Acropora monticulosa* | 0.10 | 2.68 | -0.72 | 0.43 | -1.05 | -0.22 | Both | - |
| 39 | *Acropora muricata* | 2.22 | 0.67 | -0.45 | -0.45 | 0.45 | 0.66 | Both | S,C |
| 40 | *Acropora nana* | - | -0.15 | - | 0.23 | - | 0.00 | RM | - |
| 41 | *Acropora nasuta* | 0.94 | 0.47 | -0.55 | 0.32 | -0.19 | 1.10 | Both | C |
| 42 | *Acropora paniculata* | 0.46 | 1.07 | 0.06 | 0.54 | -1.02 | 0.52 | Both | C |
| 43 | *Acropora polystoma* | - | 0.84 | - | -0.20 | - | 0.73 | RM | C |
| 44 | *Acropora robusta* | 2.32 | 0.08 | -0.26 | -0.23 | 0.74 | -0.09 | Both | - |
| 45 | *Acropora samoensis* | 0.08 | -0.31 | -0.76 | 0.51 | -0.24 | -0.57 | Both | - |
| 46 | *Acropora sarmentosa* | 2.01 | - | -0.09 | - | -0.09 | - | GBR | - |
| 47 | *Acropora secale* | 0.39 | -0.10 | -1.10 | 0.26 | -0.52 | 0.48 | Both | - |
| 48 | *Acropora selago* | 0.05 | 0.47 | -0.43 | -0.58 | -1.37 | -0.02 | Both | - |
| 49 | *Acropora solitaryensis* | 1.86 | 0.57 | 0.47 | 0.14 | 1.59 | 0.85 | Both | C |
| 50 | *Acropora speciosa* | - | 0.80 | - | -0.63 | - | -0.32 | RM | - |
| 51 | *Acropora spicifera* | - | 0.26 | - | 0.03 | - | -0.17 | RM | - |
| 52 | *Acropora striata* | - | -0.94 | - | -0.27 | - | -0.87 | RM | - |
| 53 | *Acropora subulata* | 0.80 | 0.10 | 0.62 | 0.11 | -0.39 | 0.69 | Both | C |
| 54 | *Acropora tenuis* | 0.95 | -0.29 | 0.27 | -0.56 | -0.49 | -0.37 | Both | - |
| 55 | *Acropora valenciennesi* | 2.10 | - | -0.07 | - | 0.40 | - | GBR | - |
| 56 | *Acropora valida* | -0.05 | -1.02 | -1.24 | 0.19 | -0.59 | -0.99 | Both | - |
| 57 | *Acropora vaughani* | 0.46 | 0.52 | 0.19 | 0.53 | 0.92 | 0.09 | Both | - |
| 58 | *Acropora willisae* | 0.88 | - | -1.01 | - | -0.66 | - | GBR | - |
| 59 | *Acropora yongei* | 1.07 | - | -0.79 | - | -0.61 | - | GBR | - |
| 60 | *Alveopora allingi* | 0.06 | - | 0.08 | - | -2.08 | - | GBR | - |
| 61 | *Anacropora forbesi* | - | 0.37 | - | 0.50 | - | -0.39 | RM | - |
| 62 | *Astrea curta* | 1.82 | 0.18 | 0.56 | -0.27 | 0.77 | -0.39 | Both | - |
| 63 | *Astreopora gracilis* | -2.16 | -0.60 | -1.76 | -2.51 | -0.36 | 0.40 | Both | - |
| 64 | *Astreopora listeri* | 0.77 | - | -1.55 | - | 0.82 | - | GBR | - |
| 65 | *Astreopora myriophthalma* | 0.24 | -2.58 | -1.82 | -2.27 | -0.55 | -0.79 | Both | - |
| 66 | *Astreopora ocellata* | -0.07 | - | -1.95 | - | -0.51 | - | GBR | - |
| 67 | *Australogyra zelli* | -0.28 | - | 0.84 | - | -0.29 | - | GBR | - |
| 68 | *Caulastraea furcata* | -1.14 | 2.15 | -0.18 | 0.39 | -0.37 | 0.34 | Both | - |
| 69 | *Coelastrea aspera* | 0.78 | -0.48 | 0.49 | 0.88 | -0.57 | 0.88 | Both | C |
| 70 | *Coeloseris mayeri* | 0.37 | 1.27 | -0.35 | -0.62 | -0.53 | 0.01 | Both | - |
| 71 | *Coscinaraea columna* | -1.12 | - | -3.34 | - | 0.21 | - | GBR | - |
| 72 | *Coscinaraea exesa* | -1.67 | - | -3.21 | - | -2.20 | - | GBR | - |
| 73 | *Coscinaraea monile* | - | 1.68 | - | 0.54 | - | 1.85 | RM | C |
| 74 | *Ctenactis echinata* | -0.96 | - | -4.30 | - | -0.53 | - | GBR | C |
| 75 | *Cycloseris explanulata* | - | 0.31 | - | 0.28 | - | 0.81 | RM | C |
| 76 | *Cycloseris vaughani* | - | 1.46 | - | 1.20 | - | 1.18 | RM | - |
| 77 | *Cyphastrea chalcidicum* | - | -1.61 | - | 0.06 | - | -0.02 | RM | - |
| 78 | *Cyphastrea decadia* | -0.45 | - | -1.75 | - | -1.37 | - | GBR | - |
| 79 | *Cyphastrea microphthalma* | -0.17 | -1.11 | -1.20 | -0.07 | -0.15 | -0.63 | Both | - |
| 80 | *Cyphastrea serailia* | 0.40 | - | -1.27 | - | -1.38 | - | GBR | - |
| 81 | *Diploastrea heliopora* | -0.39 | -3.14 | -2.32 | -1.80 | 0.51 | -1.74 | Both | - |
| 82 | *Dipsastraea amicorum* | 0.67 | - | 0.72 | - | 0.36 | - | GBR | - |
| 83 | *Dipsastraea danai* | 3.01 | - | 0.05 | - | 0.06 | - | GBR | - |
| 84 | *Dipsastraea favus* | 1.41 | 1.29 | 0.63 | 0.41 | -0.58 | 2.45 | Both | S,C |
| 85 | *Dipsastraea laxa* | 1.14 | -0.52 | 0.81 | 0.59 | 1.15 | -0.03 | Both | - |
| 86 | *Dipsastraea lizardensis* | 2.33 | 1.14 | 0.70 | 0.43 | 0.91 | -0.87 | Both | S,P |
| 87 | *Dipsastraea maritima* | 0.57 | -0.94 | -0.55 | 1.12 | -0.77 | -0.13 | Both | - |
| 88 | *Dipsastraea matthaii* | 1.62 | -3.19 | 0.74 | 0.71 | 1.14 | -1.66 | Both | - |
| 89 | *Dipsastraea maxima* | 0.97 | - | 0.48 | - | 0.62 | - | GBR | - |
| 90 | *Dipsastraea pallida* | 1.41 | -0.92 | 0.19 | 0.37 | 0.25 | -0.70 | Both | - |
| 91 | *Dipsastraea rosaria* | 0.54 | - | -0.29 | - | -0.99 | - | GBR | - |
| 92 | *Dipsastraea rotumana* | - | 0.23 | - | 0.62 | - | 0.77 | RM | C |
| 93 | *Dipsastraea speciosa* | 1.47 | -0.65 | 0.84 | 0.45 | -1.23 | 0.02 | Both | - |
| 94 | *Dipsastraea truncata* | 0.63 | -2.16 | -0.41 | 0.61 | -0.78 | 0.61 | Both | C |
| 95 | *Echinophyllia aspera* | -1.19 | 0.19 | -1.26 | -0.70 | -0.01 | 0.07 | Both | - |
| 96 | *Echinophyllia orpheensis* | -0.74 | 0.74 | -1.45 | -0.54 | 0.55 | -0.96 | Both | P |
| 97 | *Echinopora gemmacea* | 1.00 | -1.34 | -0.39 | 0.25 | 0.25 | -0.39 | Both | - |
| 98 | *Echinopora horrida* | 0.98 | 0.26 | -0.28 | 0.40 | 0.55 | -0.52 | Both | - |
| 99 | *Echinopora lamellosa* | 1.60 | 1.10 | -0.12 | 0.39 | -0.04 | 1.19 | Both | S,C |
| 100 | *Echinopora mammiformis* | 0.16 | - | -0.51 | - | -0.32 | - | GBR | - |
| 101 | *Euphyllia glabrescens* | - | -0.06 | - | 1.04 | - | -0.03 | RM | - |
| 102 | *Favites abdita* | 0.08 | 0.49 | 0.32 | 0.08 | 0.63 | 1.74 | Both | C |
| 103 | *Favites chinensis* | 1.00 | - | -0.60 | - | 0.72 | - | GBR | - |
| 104 | *Favites complanata* | 0.53 | -0.80 | 0.00 | -0.11 | -0.58 | 0.23 | Both | - |

| | | | | | | | | | |
|---|---|---|---|---|---|---|---|---|---|
| 105 | *Favites flexuosa* | 0.21 | -1.07 | -0.06 | 0.22 | 0.50 | -0.34 | Both | - |
| 106 | *Favites halicora* | 0.60 | -1.35 | -0.11 | 0.06 | 0.50 | 0.12 | Both | - |
| 107 | *Favites magnistellata* | -1.23 | - | -0.02 | - | 0.22 | - | GBR | - |
| 108 | *Favites melicerum* | 1.64 | - | -0.06 | - | 0.53 | - | GBR | - |
| 109 | *Favites pentagona* | 1.81 | 0.00 | 0.85 | -0.01 | 0.82 | -0.85 | Both | P |
| 110 | *Favites rotundata* | 2.18 | 0.40 | 0.07 | 0.01 | 1.03 | -1.60 | Both | P |
| 111 | *Favites valenciennesii* | 0.82 | -0.16 | -0.56 | -0.43 | 0.16 | 0.73 | Both | C |
| 112 | *Fimbriaphyllia ancora* | -0.26 | - | -1.48 | - | -0.48 | - | GBR | - |
| 113 | *Fimbriaphyllia divisa* | -1.30 | - | -1.20 | - | -0.66 | - | GBR | - |
| 114 | *Fungia fungites* | - | 1.62 | - | 1.29 | - | 0.30 | RM | - |
| 115 | *Galaxea astreata* | -0.76 | 2.26 | -1.34 | 0.87 | -1.52 | 0.33 | Both | - |
| 116 | *Galaxea fascicularis* | -0.74 | 1.73 | -0.75 | 1.18 | -1.86 | -0.06 | Both | - |
| 117 | *Galaxea horrescens* | -1.21 | - | -1.07 | - | 1.40 | - | GBR | - |
| 118 | *Gardineroseris planulata* | 0.33 | 0.23 | -1.02 | -0.76 | -0.16 | 0.12 | Both | - |
| 119 | *Goniastrea edwardsi* | 2.33 | 0.32 | 0.10 | -0.18 | 1.86 | 0.88 | Both | C |
| 120 | *Goniastrea favulus* | -0.11 | 0.15 | 0.31 | -0.06 | -0.89 | -0.29 | Both | - |
| 121 | *Goniastrea pectinata* | 1.64 | -0.34 | 0.15 | -0.12 | 0.25 | -0.95 | Both | - |
| 122 | *Goniastrea retiformis* | 3.58 | 0.46 | -0.23 | 0.08 | 1.10 | -0.83 | Both | P |
| 123 | *Goniastrea stelligera* | 1.25 | 0.11 | -0.25 | 0.49 | -0.04 | 0.48 | Both | - |
| 124 | *Halomitra pileus* | - | 2.60 | - | 1.98 | - | 1.30 | RM | C |
| 125 | *Heliofungia actiniformis* | -1.22 | - | -2.90 | - | -0.04 | - | GBR | - |
| 126 | *Herpolitha limax* | -1.35 | -2.21 | -4.22 | 1.47 | -1.58 | -1.02 | Both | - |
| 127 | *Homophyllia bowerbanki* | -0.50 | - | -2.77 | - | -0.53 | - | GBR | - |
| 128 | *Homophyllia bowerbanki* | 0.29 | - | -1.24 | - | -1.17 | - | GBR | - |
| 129 | *Hydnophora exesa* | -0.02 | 0.36 | -0.07 | -0.24 | 0.13 | 0.07 | Both | - |
| 130 | *Hydnophora microconos* | 1.82 | 0.89 | 0.26 | -0.13 | 0.96 | 0.53 | Both | S,C |
| 131 | *Hydnophora rigida* | 0.89 | - | 0.15 | - | 0.60 | - | GBR | - |
| 132 | *Isopora brueggemanni* | 2.12 | - | -0.56 | - | 0.17 | - | GBR | - |
| 133 | *Isopora cuneata* | 1.64 | - | -0.53 | - | -0.26 | - | GBR | - |
| 134 | *Isopora palifera* | 0.96 | 0.94 | -0.79 | 1.53 | 0.15 | -0.08 | Both | - |
| 135 | *Leptastrea inaequalis* | -1.98 | -1.24 | -3.02 | 0.64 | -0.76 | -0.75 | Both | - |
| 136 | *Leptastrea purpurea* | -0.80 | 0.82 | -2.91 | 0.34 | 0.39 | -0.05 | Both | - |
| 137 | *Leptastrea purpurea* | -1.37 | -0.46 | -2.81 | 0.34 | -1.05 | 0.02 | Both | - |
| 138 | *Leptastrea transversa* | -1.12 | -0.69 | -2.94 | 0.83 | -1.72 | 0.09 | Both | - |
| 139 | *Leptoria phrygia* | 2.90 | 0.32 | 1.12 | 0.44 | 1.47 | -0.69 | Both | P |
| 140 | *Leptoseris hawaiiensis* | - | -1.13 | - | 0.19 | - | -0.89 | RM | - |
| 141 | *Leptoseris mycetoseroides* | 0.66 | 1.13 | -0.70 | 0.45 | -0.80 | -0.04 | Both | - |
| 142 | *Leptoseris scabra* | - | 0.66 | - | 0.12 | - | 1.21 | RM | C |
| 143 | *Leptoseris yabei* | 0.67 | -0.22 | -0.68 | 0.46 | 1.35 | 1.33 | Both | C |
| 144 | *Lithophyllon concinna* | -1.47 | - | -2.81 | - | -0.37 | - | GBR | - |
| 145 | *Lobophyllia agaricia* | -0.46 | - | -1.29 | - | -0.08 | - | GBR | - |
| 146 | *Lobophyllia corymbosa* | 0.43 | 1.12 | -1.43 | 0.93 | -0.42 | 0.74 | Both | C |
| 147 | *Lobophyllia flabelliformis* | -0.27 | - | -1.40 | - | 0.16 | - | GBR | - |
| 148 | *Lobophyllia hataii* | -0.86 | - | -1.68 | - | -0.10 | - | GBR | - |
| 149 | *Lobophyllia hemprichii* | 0.22 | - | -1.74 | - | 0.42 | - | GBR | - |
| 150 | *Lobophyllia radians* | -1.05 | - | -1.31 | - | -1.65 | - | GBR | - |
| 151 | *Lobophyllia recta* | 0.87 | 0.86 | -1.14 | 0.19 | 1.26 | -0.14 | Both | - |
| 152 | *Lobophyllia robusta* | -1.71 | - | -1.38 | - | 0.10 | - | GBR | - |
| 153 | *Lobophyllia valenciennesii* | -0.50 | - | -1.32 | - | -0.12 | - | GBR | - |
| 154 | *Lobophyllia vitiensis* | -1.16 | - | -1.48 | - | 0.31 | - | GBR | - |
| 155 | *Merulina ampliata* | 0.07 | 0.80 | 0.13 | 0.22 | -0.76 | 0.26 | Both | - |
| 156 | *Merulina scabricula* | 1.76 | - | -0.02 | - | -0.37 | - | GBR | - |
| 157 | *Micromussa lordhowensis* | -1.42 | - | -2.50 | - | -0.36 | - | GBR | - |
| 158 | *Moseleya latistellata* | -2.06 | - | -0.36 | - | 0.69 | - | GBR | - |
| 159 | *Mycedium elephantotus* | -0.06 | 1.29 | -0.89 | 0.05 | -0.06 | 0.45 | Both | - |
| 160 | *Oulophyllia bennettae* | 1.32 | - | 0.12 | - | 0.17 | - | GBR | - |
| 161 | *Oulophyllia crispa* | 0.43 | 0.91 | -0.03 | 1.01 | 0.46 | 0.13 | Both | - |
| 162 | *Oxypora lacera* | 0.67 | - | -1.65 | - | 0.19 | - | GBR | - |
| 163 | *Pachyseris rugosa* | -0.83 | -1.13 | -0.95 | -0.68 | -0.39 | -0.94 | Both | - |
| 164 | *Pachyseris speciosa* | 2.03 | -0.15 | -1.70 | -0.49 | 0.46 | 0.50 | Both | C |
| 165 | *Palauastrea ramosa* | 1.68 | - | -0.64 | - | -0.10 | - | GBR | - |
| 166 | *Paragoniastrea australensis* | 0.53 | - | 0.21 | - | -0.48 | - | GBR | - |
| 167 | *Paragoniastrea russelli* | 1.02 | -0.46 | 0.33 | 0.29 | 1.22 | -0.58 | Both | - |
| 168 | *Pavona clavus* | - | -0.74 | - | -0.88 | - | 1.32 | RM | C |
| 169 | *Pavona decussata* | -0.39 | - | -0.88 | - | -1.42 | - | GBR | - |
| 170 | *Pavona duerdeni* | -0.11 | - | -0.97 | - | 0.13 | - | GBR | - |
| 171 | *Pavona explanulata* | - | -3.01 | - | -0.70 | - | -1.10 | RM | - |
| 172 | *Pavona maldivensis* | -1.55 | -0.95 | -1.18 | -0.74 | -0.98 | 1.10 | Both | C |
| 173 | *Pavona varians* | -2.09 | -2.31 | -1.09 | -1.50 | 0.19 | -1.99 | Both | - |

| | | | | | | | | | |
|---|---|---|---|---|---|---|---|---|---|
| 174 | *Pavona venosa* | - | -1.92 | - | -0.71 | - | -1.08 | RM | - |
| 175 | *Pectinia alcicornis* | - | -0.10 | - | 0.18 | - | -0.32 | RM | - |
| 176 | *Pectinia lactuca* | -2.38 | -0.25 | -0.36 | 0.09 | 0.40 | 1.07 | Both | C |
| 177 | *Pectinia paeonia* | 0.34 | - | -0.78 | - | 2.00 | - | GBR | - |
| 178 | *Physogyra lichtensteini* | -2.42 | -2.06 | -1.82 | -1.35 | -1.48 | -1.10 | Both | - |
| 179 | *Platygyra acuta* | 0.19 | - | 0.10 | - | -0.86 | - | GBR | - |
| 180 | *Platygyra contorta* | 0.01 | - | 0.36 | - | -0.50 | - | GBR | - |
| 181 | *Platygyra daedalea* | 2.41 | 0.10 | 0.84 | 0.31 | 0.58 | 0.33 | Both | - |
| 182 | *Platygyra lamellina* | 1.33 | 1.92 | 0.45 | -0.41 | 0.74 | 0.46 | Both | - |
| 183 | *Platygyra pini* | 1.36 | -0.56 | 0.69 | -0.02 | 0.77 | -0.99 | Both | - |
| 184 | *Platygyra ryukyuensis* | 0.90 | - | 0.21 | - | 0.08 | - | GBR | - |
| 185 | *Platygyra sinensis* | 1.44 | - | 0.78 | - | 0.38 | - | GBR | - |
| 186 | *Platygyra verweyi* | 2.05 | - | 0.68 | - | 0.97 | - | GBR | - |
| 187 | *Plerogyra sinuosa* | - | -3.55 | - | -1.17 | - | 0.58 | RM | C |
| 188 | *Plesiastrea versipora* | -1.10 | -1.45 | -3.34 | -0.18 | 0.21 | -0.52 | Both | - |
| 189 | *Pleuractis paumotensis* | -2.16 | - | -3.64 | - | -0.07 | - | GBR | - |
| 190 | *Pocillopora damicornis* | 2.72 | -2.89 | 0.77 | 0.01 | 1.13 | 0.69 | Both | C |
| 191 | *Pocillopora grandis* | 1.48 | 1.43 | 0.50 | 0.14 | 0.63 | 2.30 | Both | S,C |
| 192 | *Pocillopora indiania* | - | 0.30 | - | 1.51 | - | 0.31 | RM | - |
| 193 | *Pocillopora meandrina* | - | -0.88 | - | 0.43 | - | 0.45 | RM | - |
| 194 | *Pocillopora verrucosa* | 3.31 | - | 0.68 | - | 0.57 | - | GBR | - |
| 195 | *Pocillopora verrucosa* | 1.83 | -1.31 | 0.97 | -0.01 | -0.19 | -0.08 | Both | - |
| 196 | *Podabacia crustacea* | -0.85 | - | -3.82 | - | 0.09 | - | GBR | - |
| 197 | *Polyphyllia talpina* | -1.80 | - | -3.96 | - | -0.49 | - | GBR | - |
| 198 | *Porites cylindrica* | - | -1.04 | - | -3.94 | - | 1.80 | RM | C |
| 199 | *Porites lichen* | 0.74 | -3.39 | -0.12 | -3.90 | 0.84 | -1.02 | Both | - |
| 200 | *Porites rus* | - | -3.39 | - | -3.78 | - | -0.74 | RM | - |
| 201 | *Psammocora digitata* | - | -4.63 | - | -0.18 | - | -0.83 | RM | - |
| 202 | *Psammocora profundacella* | - | -2.08 | - | -0.82 | - | -0.73 | RM | - |
| 203 | *Sandalolitha robusta* | -1.39 | - | -3.60 | - | 0.09 | - | GBR | - |
| 204 | *Scapophyllia cylindrica* | 0.62 | - | 0.23 | - | 0.16 | - | GBR | - |
| 205 | *Seriatopora hystrix* | 1.66 | - | 1.71 | - | -1.27 | - | GBR | - |
| 206 | *Siderastrea savignyana* | - | -1.19 | - | -0.95 | - | 0.33 | RM | - |
| 207 | *Stylocoeniella guentheri* | -0.20 | -1.46 | -0.95 | -0.52 | -0.62 | 1.29 | Both | C |
| 208 | *Stylophora pistillata* | 3.25 | - | 2.44 | - | 1.06 | - | GBR | - |
| 209 | *Turbinaria frondens* | -1.62 | - | -2.50 | - | -0.43 | - | GBR | - |
| 210 | *Turbinaria mesenterina* | -1.93 | -0.20 | -2.16 | 0.24 | -1.34 | -0.70 | Both | P |
| 211 | *Turbinaria patula* | -1.27 | - | -2.96 | - | -0.33 | - | GBR | - |
| 212 | *Turbinaria peltata* | -1.69 | - | -2.52 | - | -1.01 | - | GBR | - |
| 213 | *Turbinaria reniformis* | -0.55 | - | -2.51 | - | 0.12 | - | GBR | - |
| 214 | *Turbinaria stellulata* | -1.07 | 0.76 | -1.82 | 0.43 | -0.57 | 0.01 | Both | - |

**Table S3** Posterior medians for each species derived from the two phylogenetic linear mixed model analyses. GBR: Great Barrier Reef, RM: Republic of Maldives, Total: overall bleaching response, Phylogen: phylogenetically heritable component, Contemp: contemporary component, S: highest quartile of susceptibility in both regions, C: highest quartile for contemporary effects in the RM, P: lowest quartile for contemporary effects and highest two quartiles for susceptibility in RM.

| Region | Proportion of Variation Explained | | |
|---|---|---|---|
| | **Phylogenetic** | **Contemporary** | **Local-Scale** |
| Maldives | 0.261 (0.212,0.306) | 0.168 (0.142, 0.189) | 0.035 (0.028,0.036) |
| GBR | 0.467 (0.419, 0.498) | 0.048 (0.028,0.049) | 0.012 (0.070,0.106) |

**Table S4** Phylogenetic mixed model analyses exploring the phylogenetic uncertainty in the posterior median estimates. Presented are the 0.025 and 0.975 quantiles of the distribution of posterior medians calculated from each of the 1000 posterior samples of the super-tree (Huang & Roy, 2016). The proportion of the variation in bleaching explained by phylogenetically heritable (phylogenetic), recent adaptation/ acclimatization (contemporary) and local-scale effects.

| Region | Proportion of Variation Explained | | |
|---|---|---|---|
| | **Phylogenetic** | **Contemporary** | **Local-Scale** |
| Maldives | 0.370 (0.040,0.639) | 0.047 (0.048, 0.266) | 0.032 (0.012,0.100) |
| GBR | 0.455 (0.280, 0.661) | 0.071 (0.014,0.101) | 0.001 (0.001,0.042) |

**Table S5** Phylogenetic mixed model analyses including groups of species of the genera *Porites, Montipora, Goniopora* and *Cycloceris* that were not possible to identify to species. The proportion of the variation in bleaching explained by phylogenetically heritable (phylogenetic), recent adaptation/ acclimatization (contemporary) and local-scale effects.

| Region | Proportion of Variation Explained | | |
|---|---|---|---|
| | Phylogenetic | Contemporary | Local-Scale |
| Maldives | 0.416 (0.076,0.692) | 0.088 (0.034, 0.243) | 0.036 (0.014,0.112) |
| GBR | 0.496 (0.295, 0.746) | 0.035 (0.008,0.098) | 0.014 (0.002,0.075) |

**Table S6** Phylogenetic mixed model analyses using a phylogeny based only on molecular data (Huang & Roy, 2016). The proportion of the variation in bleaching explained by phylogenetically heritable (phylogenetic), recent adaptation/ acclimatization (contemporary) and local-scale effects.

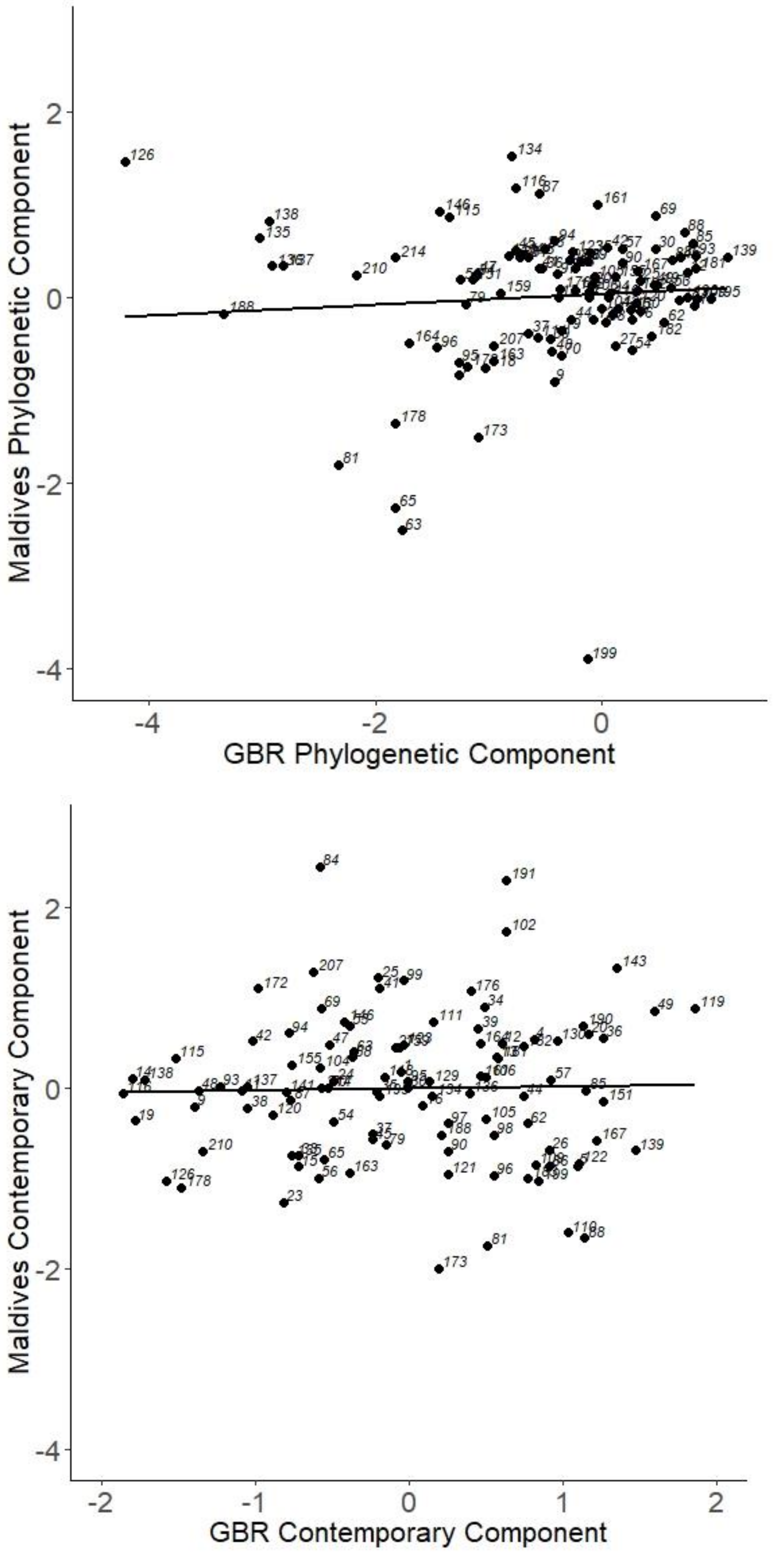


**Figure S1.** Comparison between regions for (a) the phylogenetic component of species' bleaching response and (b) contemporary component. Labels denote species number given in Table S2. Responses are derived the species posterior medians derived from phylogenetic mixed model analyses. Lines denote correlation.

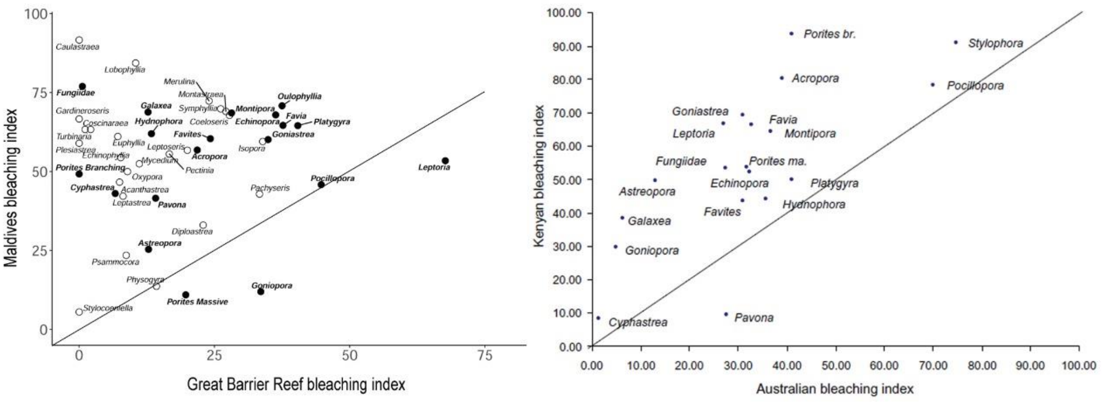


**Figure S2.** Comparison of results from (a) the present study and (b) those reported by a study of regional variation (McClanahan et al., 2004) using a bleaching mortality index (BMI) for species pooled into genera. Open symbols and a lighter label denote additional genera not reported by McClanahan et al. (2004). Genus *Stylophora* was not detected in the Maldives surveys, but was reported until recently and was likely highly susceptible (Muir et al., 2017).
